# Genetic interference with HvNotch provides new insights into the role of the Notch-signalling pathway for developmental pattern formation in *Hydra*

**DOI:** 10.1101/2024.01.31.578171

**Authors:** Qin Pan, Moritz Mercker, Alexander Klimovich, Jörg Wittlieb, Anna Marciniak-Czochra, Angelika Böttger

## Abstract

The Notch-signalling pathway plays an important role in pattern formation in *Hydra*. Using pharmacological Notch inhibitors (DAPT and SAHM1), it has been demonstrated that HvNotch is required for head regeneration and tentacle patterning in *Hydra*. HvNotch is also involved in establishing the parent-bud boundary and instructing buds to develop feet and detach from the parent. To further investigate the functions of HvNotch, we successfully constructed NICD (HvNotch intracellular domain)-overexpressing and HvNotch-knockdown transgenic *Hydra* strains. NICD-overexpressing transgenic *Hydra* showed a pronounced inhibition on the expression of predicted HvNotch-target genes, suggesting a dominant negative effect of ectopic NICD. This resulted in a “Y-shaped” phenotype, which arises from the parent-bud boundary defect seen in polyps treated with DAPT. Additionally, “multiple heads”, “two-headed” and “ectopic tentacles” phenotypes were observed. The HvNotch-knockdown transgenic *Hydra* with reduced expression of HvNotch exhibited similar, but not identical phenotypes, with the addition of a “two feet” phenotype. Furthermore, approximately 20% of the HvNotch-knockdown polyps were unable to regenerate a new head after decapitation. We integrated these findings into a mathematical model based on long-range gradients of signalling molecules underlying sharply defined positions of HvNotch-signalling cells at the *Hydra* tentacle and bud boundaries.

## Introduction

The freshwater polyp *Hydra* (Cnidaria) has a simple body plan, comprising a single axis with a hypostome surrounded by a ring of tentacles at the oral end and a peduncle with a basal disk at the aboral end. *Hydra* can reproduce asexually by budding. The entire body consists of three cell lineages: ectodermal epithelial cells, endodermal epithelial cells and interstitial cells. The epithelial cells represent two self-renewing epithelia with continuous proliferation in the body column. At the tentacle boundaries, these cells undergo mitotic exit and differentiate into battery cells, while at the aboral end, they differentiate into peduncle cells^1^. The interstitial cell lineage is located in the spaces between the epithelial cells and consists of multipotent stem cells and their differentiation products, including nematocytes, nerve cells, gland cells, and germ cells^2^.

Due to its ongoing self-renewal, the adult *Hydra* harbours all the necessary information for body patterning, enabling an almost unlimited capacity of regenerating lost body parts. In 1909, Ethel Browne performed grafting experiments that demonstrated the ability of *Hydra* head tissue to induce the formation of a new hydranths when transplanted into the body column of recipient polyps. This process involved recruiting recipient tissue into the new head structures and indicated the presence of an “organiser” function of these tissues, a term created by Hans Spemann and Hilde Mangold only in 1923 to describe a tissue in amphibian embryos with similar abilities^3,4^. Further transplantation studies revealed that the head-forming potential of transplants gradually decreases with their distance from the head of the donor animal but increases when positioned further away from the head in the host animal^5,6^. These data were interpreted according to a reaction-diffusion model developed by Gierer and Meinhardt in 1972 with two major assumptions: (1) the head organizer produces a self-activating head activation signal (HA) with a short range and a long-range inhibition signal (HI). Both signals exist in a gradient pattern from the head to the body column^5-10^; (2) A head activation gradient is present throughout the whole length of the body column of *Hydra*^11^.This gradient serves as a slowly changing long-term storage of the body axis gradients and plays a crucial role in the interplay between different pattern formation systems^12^.

It has been suggested that canonical Wnt-signalling plays a major role in head activation and that nuclear β–catenin defines the head activation gradient along the *Hydra* body axis^13,14^. However, ectopic activation of Wnt-signalling using the GSK-3 inhibitor alsterpaullone led to the formation of ectopic tentacles instead of complete ectopic heads, indicating some missing links.

The Notch signalling pathway plays a crucial role in cell-fate determination and pattern formation by regulating cell-to-cell communication during development. The Notch receptor and its ligands are both transmembrane proteins. Ligands in one cell trans-activate Notch in a neighbouring cell, inducing two proteolytic cleavages to release the intracellular domain of Notch (NICD) from the membrane to the nucleus. NICD then binds to transcriptional regulators of the CSL-family (CBF1, Suppressor of Hairless, Lag2) and co-activates transcriptional targets^15,16^.

Additionally, ligand and receptor in the same cell mutually inhibit one another^17-19^. Cells with a high ligand concentration and a low Notch concentration are in a preferentially sending state. Conversely, cells are in a receiver state when the concentration of Notch-receptor is higher than that of ligand. Thus, two mutually exclusive states of Notch activation are created. The activity of Notch reaches its peak when these two states are next to each other, then inducing the formation of a sharp tissue border, which may explain many of the observed tissue patterning functions of Notch-signalling^19,20^.

The Notch protein (HvNotch), a ligand (HyJagged) and the canonical signal transduction pathway are conserved in *Hydra*. The *Hydra* Hes-family bHLH transcription factor 2 (HyHes) has been shown to be a target for transcriptional activation by NICD^21,22^. The Notch signalling pathway in *Hydra* plays a critical role in regulating tentacle boundaries and head regeneration after decapitation. Blocking Notch signalling with DAPT in adult *Hydra* leads to the formation of abnormal heads with irregularly arranged tentacles^23^. The Notch signalling pathway is also essential for the formation of the parent-bud boundary^23,24^. When Notch-signalling is inhibited at the parent-bud boundary, the buds fail to form a foot and remain attached to the parent, resulting in the formation of Y-animals. Through a differential gene regulation analysis with Notch-inhibited *Hydra*, the transcriptional repressor HyHes, the putative transcriptional repressor of Wnt3, Sp5^25^ and the tentacle boundary gene HyAlx^26^ were identified as potential transcriptional target genes for NICD. Moreover, the *Hydra* Fos-homolog kayak was found to be up-regulated after DAPT inhibition, indicating that it could be a potential target of Notch-induced transcriptional repressors, such as HyHes^27^.

Previous insights into these Notch-functions had been obtained by using pharmacological inhibitors, DAPT or SAHM1. However, drug treatment was always only sustained for 48 h, making it impossible to observe long-term effects of Notch-ablation^23^. Additionally, it is important to consider the potential side effects of using pharmacological drugs in animals. Therefore, to further understand the function of the Notch signalling pathway in *Hydra*, an alternative approach involving genetic interference with HvNotch was considered.

Here we created Notch transgenic *Hydra* strains, one overexpressing NICD in either ectodermal or endodermal epithelial cells, and another expressing an interfering HvNotch-hairpin-RNA mediating Notch-knockdown in both epithelial cell layers. We monitored these strains over extended periods of time and compared the phenotypes observed in ectodermal and endodermal NICD-overexpressing polyps and in HvNotch-knockdown polyps. We found similar phenotypes as had been observed after inhibition with DAPT or SAHM1, confirming that HvNotch functions at tissue boundaries. Moreover, we obtained evidence for an additional function of the Notch-signalling pathway in regulating the head activation gradient along the *Hydra* body axis. Finally, we provided an initial mathematic model to explain how HvNotch functions to ensure spatio-temporal timing of Notch-signalling at the parent-bud boundary.

## Results

### NICD-overexpressing transgenic *Hydra*

#### The establishment of NICD (Notch intracellular domain)-overexpressing transgenic *Hydra*

To create NICD-overexpressing *Hydra*, we cloned the NICD encoding segment of HvNotch into the pHyVec11 vector, which also contains a downstream DsRed fluorescence (Fig. 1A). After injecting this plasmid into *Hydra* embryos, we obtained 60 embryos with NICD-pHyVec11 injection and 47 embryos with control-pHyVec11 injection. Only one polyp (named 4# strain) exhibited obvious DsRed signals in the NICD-pHyVec11 group, in comparison to the control with 14 DsRed-positive polyps (supplementary Fig.S1A and Fig.S1B), suggesting a negative effect of NICD-overexpression on embryogenesis. Through the selection of buds with enriched transgenic cell pools and regeneration experiments, we obtained uniformly transgenic *Hydra* strains with NICD-overexpression in the entire ectoderm or in the entire endoderm (Fig.1B). Both of these transgenic strains displayed more than 10-fold higher expression of NICD at the RNA level as measured by RT-qPCR. RT-qPCR using primers for detecting full length HvNotch transcripts indicated that expression of the endogenous HvNotch was unaffected (Fig.1C).

**Fig. 1.**
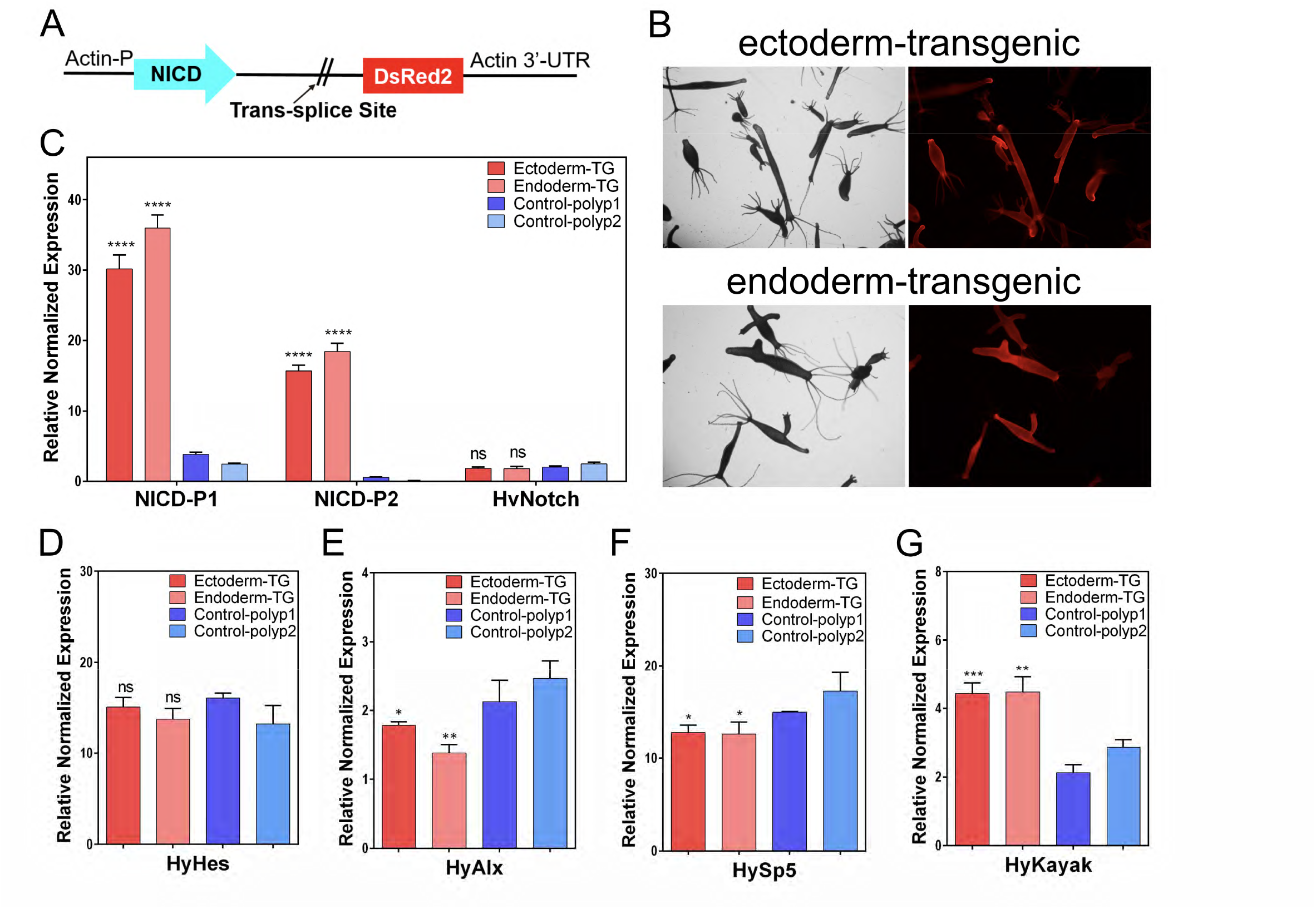
HvNICD-overexpressing transgenic *Hydra* and the expression of Notch-target genes. (A) Vector pHyVec11 with HvNICD-insert; HvNICD sequence is under the control of the *Hydra*-actin-promoter. The DsRed-sequence is included in an operon with NICD and expressed independently of HvNICD after trans-splice leader addition. (B) Fully transgenic *Hydra* expressing the transgenes in the whole ectoderm (upper panels, referred to as Ectoderm-TG), and in the whole endoderm (lower panels, referred to as Endoderm-TG). (C) Diagram presents the relative normalized expression of HvNotch and HvNICD, as determined by RT-qPCR with mRNA from Ectoderm-TG and Endoderm-TG polyps in comparison with control groups (control-polyp1 and control-polyp2); the p-values related to the average values of control polyps 1 and 2 for ectoderm-TG: p < 0.0001 and endoderm-TG: p < 0.0001. Primers for NICD-P1 and NICD-P2 were designed for amplification of HvNICD-sequence, primers for HvNotch were designed for amplification of Notch-extracellular domain sequence. (D-G) Diagram presents the relative normalized expression of HvNotch-target genes. (D) HyHes did not show significant difference between Ectoderm-TG /Endoderm-TG and the average of two control groups. (E) HyAlx with significant downregulation in Ectoderm-TG (p = 0.013) and Endoderm-TG (p = 0.0025). (F) HySp5 with downregulation in both Ectoderm-TG (p = 0.012) and Endoderm-TG (p = 0.022). (G) HyKayak with significant up-regulation in Ectoderm-TG (p = 0.0009) and Endoderm-TG (p = 0.002); p-values always related to the average of both control groups.

### The expression level of target genes in NICD-overexpressing transgenic *Hydra*

RT-qPCR analysis revealed that the predicted Notch-target genes HyAlx and HySp5 both showed a significant downregulation (Fig.1E, 1F), whereas HyHes was not significantly affected in NICD-overexpressing polyps (Fig.1D). Moreover, the expression of the *Hydra* Fos-homolog kayak (HyKayak) was clearly higher in both NICD-overexpressing strains in comparison with controls (Fig. 1G). These gene expression analyses indicate that NICD-overexpression leads to similar effects on the expression of potential Notch-target genes, as observed with DAPT-treatment (HyAlx and HySp5 down, Hykayak up, see ^27^). This suggests that NICD-overexpression has a dominant negative effect on the transcriptional activity of HvNotch and equals loss-of-function mutants.

### Phenotypes of NICD-overexpressing transgenic *Hydra*

At initial stages of obtaining fully transgenic *Hydra*, we detected “ectopic tentacles” and “two-headed” phenotypes. Approximately 10% of the polyps overexpressing NICD displayed one or two additional tentacles in the body column (Fig.2A). This phenomenon was observed in both ectodermal (Fig.2A, a-b) and endodermal transgenic polyps (Fig.2A, c-d) (referred to as Ectoderm-TG and Endoderm-TG). Furthermore, 5% of Ectoderm-TG polyps exhibited an ectopic head along the body column, which occurred near the budding region (Fig.2B, a-b), in the middle of the body column (Fig.2B, c) or at the apical end (Fig.2B, d). These phenotypes are similar to those induced with the GSK3-β inhibitor alsterpaullone (ectopic tentacles), transgenic *Hydra* overexpressing stabilised β-catenin (multiple heads along the body column), or Sp5-RNAi transgenic animals (“bouquet” like with two heads at the apical end)^14,28,29^.

After three weeks, in addition to the “ectopic tentacles” and “two-headed” phenotypes (supplementary Table S1 and Fig.S2), around 4% of the polyps presented multiple heads in the budding region, with a maximum of five ectopic heads (Fig.2C, a-c: ectoderm; d: endoderm). This phenomenon was similar to the “bouquet”-like animals, however, in our strains the “bouquet” seemed to appear only on developing buds and not on the parent head. Moreover, the majority of these ectopic heads presented normal hypostomes, while a few displayed incomplete head structures.

Additionally, we observed the presence of “Y-shaped” polyps, which closely resembled the phenotypes of DAPT-treated Hydra^24^ (Fig.2D, a-c: ectoderm; d: endoderm). However, most of these Y-shaped polyps generated new buds that detached from their parent in a normal manner during the subsequent culture process, suggesting that NICD-overexpression only hindered the detachment of buds at specific time points. We also observed more complex phenotypes, combining at least two of the aforementioned phenotypes. These included “Y-shaped” animals with “two-headed” (Fig.2E, a-b), “Y-shaped” polyps with “ectopic tentacles” (Fig. 2E, b-d) and animals with dual “Y-shaped” features probably resulting from repeatedly disrupted bud detachment (Fig.2E, d).

**Fig. 2.**
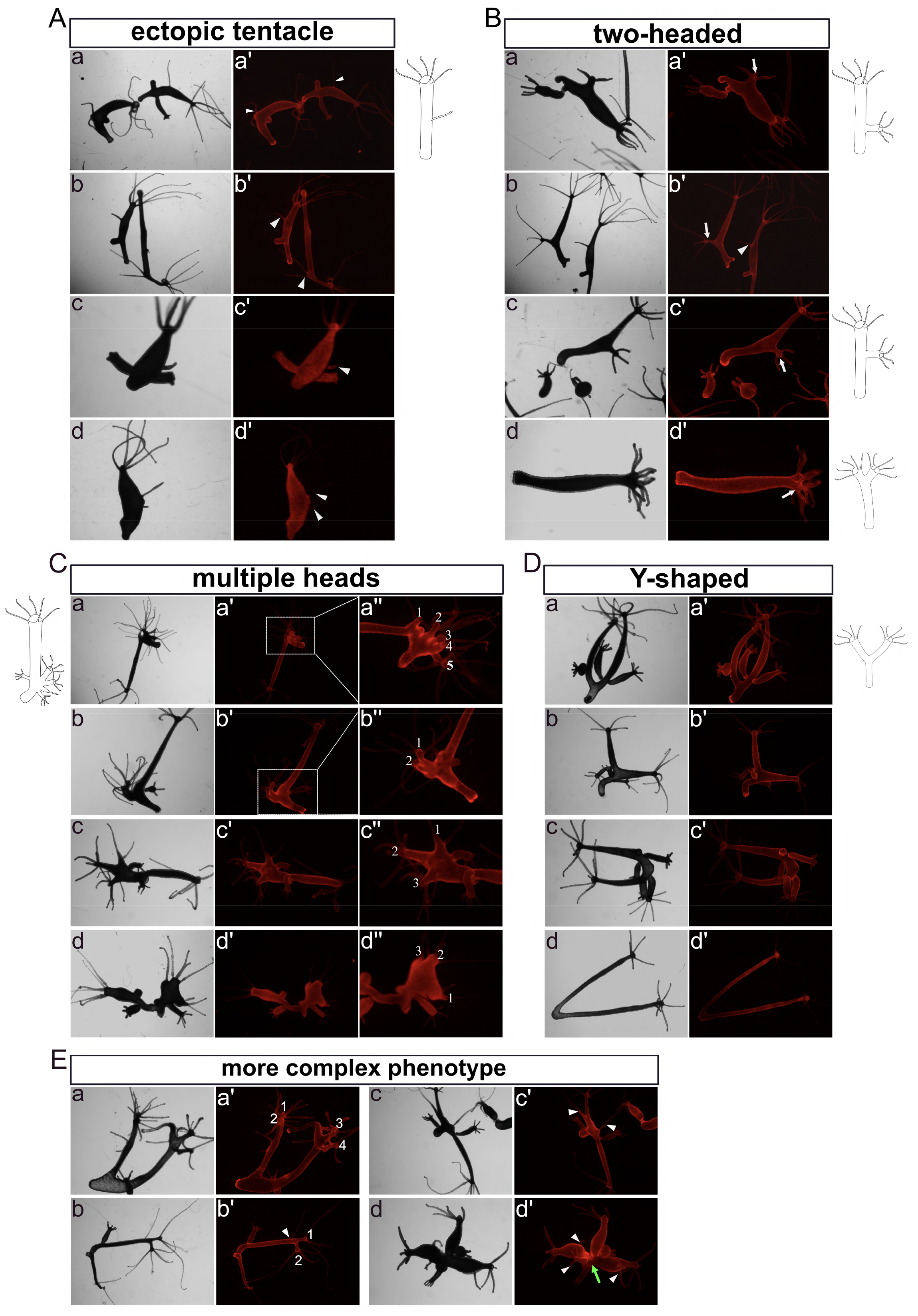
Images and drawings of Phenotypes observed in fully HvNICD-overexpressing transgenic *Hydra*. The images labeled with a-d represent light-microscopy, while a’-d’ display Ds-Red fluorescence. (A, B) Phenotypes observed in the initial stage included “ectopic tentacles” (A, a-b Ectoderm-TG; c-d Endoderm-TG) and “two-headed” (B, all Ectoderm-TG). (C, D, E) After three weeks, the observed phenotypes included “multiple heads” (C, a-c Ectoderm-TG; d Endoderm-TG; the ectopic heads are numbered), “Y-shaped animals” (D, a-c Ectodermal TG; d Endodermal TG) and combined phenotypes (E, a-d Ectoderm TG). Ectopic tentacles are indicated by white triangles, ectopic heads are indicated by white arrows.

During the following month, these distinctive phenotypes remained observable (supplementary Table S1). The number of ectopic tentacles slightly increased, with some ectoderm-TG polyps now possessing more than three in the body column or in the foot region (Fig.3A). The “multi-headed” phenotype became less intricate, with fewer extra heads/axes compared to the second stage (Fig. 3B, a: ectoderm; b: endoderm). “Two-headed” and “Y-shaped” polyps remained (Fig.3C, 3D). It is worth noting that a new type of “Y-shaped” polyps emerged with a shared head and a shared foot (Fig.3D, a: ectoderm; b: endoderm).

**Fig. 3.**
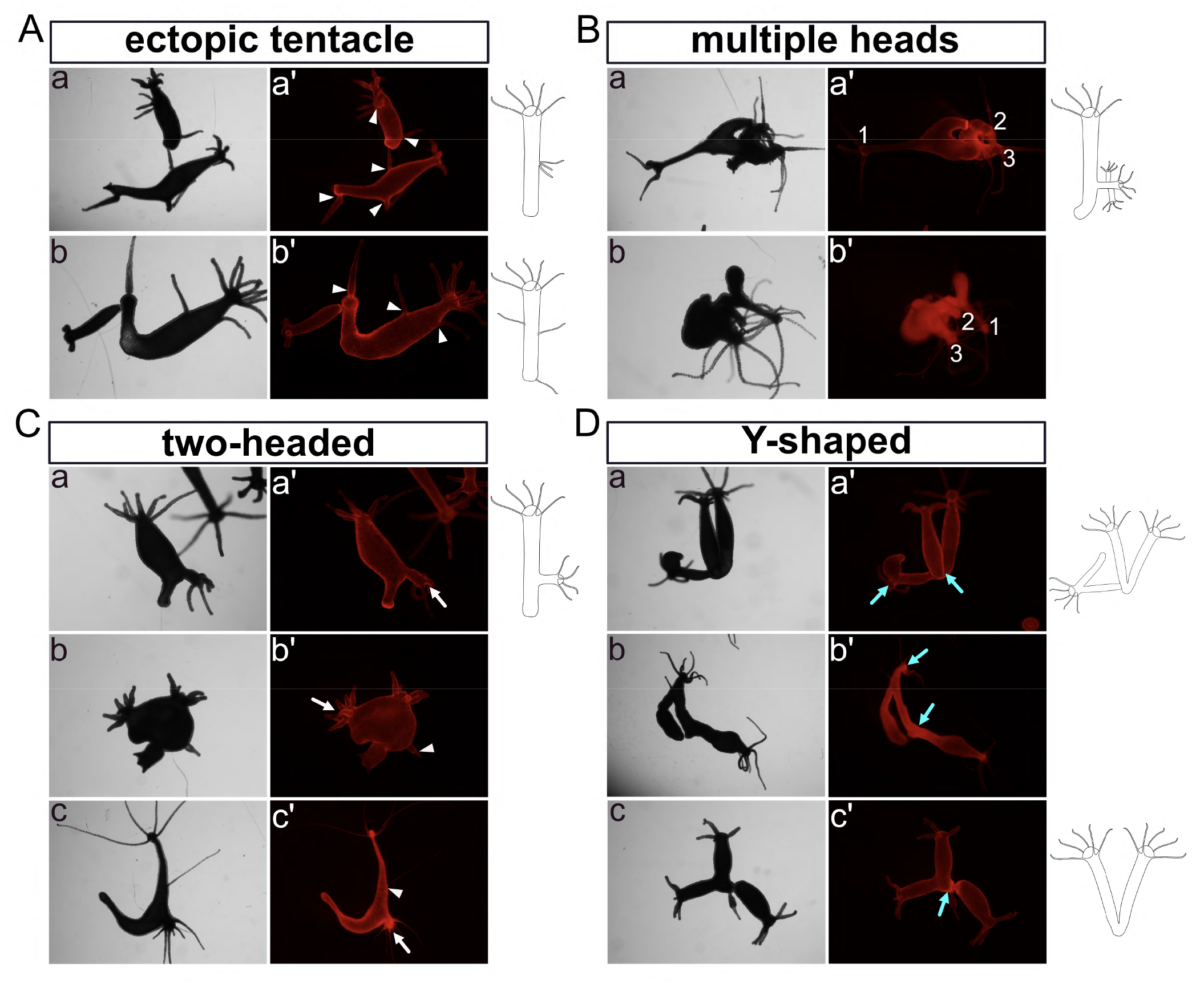
Images and drawings of phenotypes observed in long–term sustained HvNICD-overexpressing transgenic *Hydra*. The images labeled with a-c represent light-microscopy, while a’-c’ display Ds-Red fluorescence. (A) “Ectopic tentacle” phenotype (a-b Ectoderm-TG; with ectopic tentacles indicated by white triangles). (B) “Multiple heads” with less ectopic heads compared to earlier stages (a Ectoderm-TG; b Endoderm-TG; the ectopic heads are numbered). (C) “two-headed” polyps with extra heads indicated by white arrows located in the lower part of the body column (a-b Ectoderm-TG; c endoderm-TG). (D) “Y-shaped” animals characterized by either shared feet or a shared head, which are indicated by blue arrows (a-c, Ectoderm TG; b Endoderm-TG).

### NICD-overexpressing *Hydra* had a normal head regeneration process

We proceeded with a head regeneration experiment by removing heads just underneath the tentacle ring using *Hydra* that overexpressed NICD but had normal axis pattering. The NICD-overexpressing polyps did not display significant differences in their abilities to regenerate, including regeneration time and patterns of the regenerated heads, in comparison to control group (supplementary Fig.S3A: 24 h after regeneration; S3B: 72 h after regeneration).

### Notch-Knockdown transgenic Hydra

#### The establishment of Notch-knockdown transgenic *Hydra*

In order to generate Notch-knockdown *Hydra*, a sequence of the HvNotch-receptor gene (nucleotide 1763-2283) was cloned in both sense and antisense directions into the pHyVec12 vector to be transcribed into a hairpin RNA. In this vector, the *Hydra* actin promoter controls the expression of the HvNotch-hairpin, and an internal sequence allows for the addition of a splice leader between the hairpin and downstream DsRed sequences (Fig.4A). Consequently, two independent transcripts can be produced: DsRed2-mRNA and Notch-hairpin-RNA^30^.

After microinjecting the plasmid into embryos, a total of 9 polyps displaying mosaic signals developed in the HvNotch-knockdown group and 10 polyps in the control group injected with control-pHyVec12, which might indicate that Notch-hairpin expression did not have an effect on embryogenesis (supplementary Fig.S4A). Through a series of selection processes, we obtained 4 transgenic strains, in which Notch-knockdown occurred throughout the ectoderm and endoderm (supplementary Fig.S4A, S4B). All of these transgenic strains exhibited a significant decrease of approximately 3-fold in HvNotch expression at the mRNA level compared to the control groups, which included 3 strains injected with control-pHyVec12 and 1 strain of polyps injected with Notch-hairpin pHyVec12, but without any DsRed expression (referred to as empty polyps) (Fig.4B). RT-qPCR did not reveal substantial differences in the expression of potential HvNotch-target genes, as shown for HyHes, HyAlx, HySp5 and Hykayak (supplementary Fig. S4C).

### Phenotypes of Notch-knockdown transgenic *Hydra*

Two weeks after obtaining fully transgenic epithelial HvNotch-knockdown strains, we found “Y-shaped” polyps in the 11# strain. The buds developed in the lower part of the body column and then moved into the foot area within a week (Fig.4C). There was one case of two-headed polyp, in which the ectopic head initially developed in the lower part of the body column but then moved to the foot area within a week (supplementary Fig.S5A). Unfortunately, this polyp was unable to catch food and subsequently died. We noticed that dying polyps generally had very strong DsRed signals. However, immunofluorescence staining of Cadherin and acetylated Tubulin indicated that this polyp possessed normal neural nets and nematocyte-capsules (supplementary Fig.S5B).

During the subsequent three months, additional phenotypes began to manifest in all 4 strains of Notch-knockdown polyps (supplementary Table S2). We observed the presence of “ectopic tentacles” on the body column (Fig.5A), which closely resembled those observed in NICD-overexpressing polyps (Fig.5A, a). However, it is worth noting that most of these ectopic tentacles showed thickening of different lengths at their bases (Fig. 4C, a and Fig.5A, b-e), similar to the irregular head structures previously observed in *Hydra* heads and heads of non-detached buds after DAPT-treatment^23,24^.

**Fig. 4.**
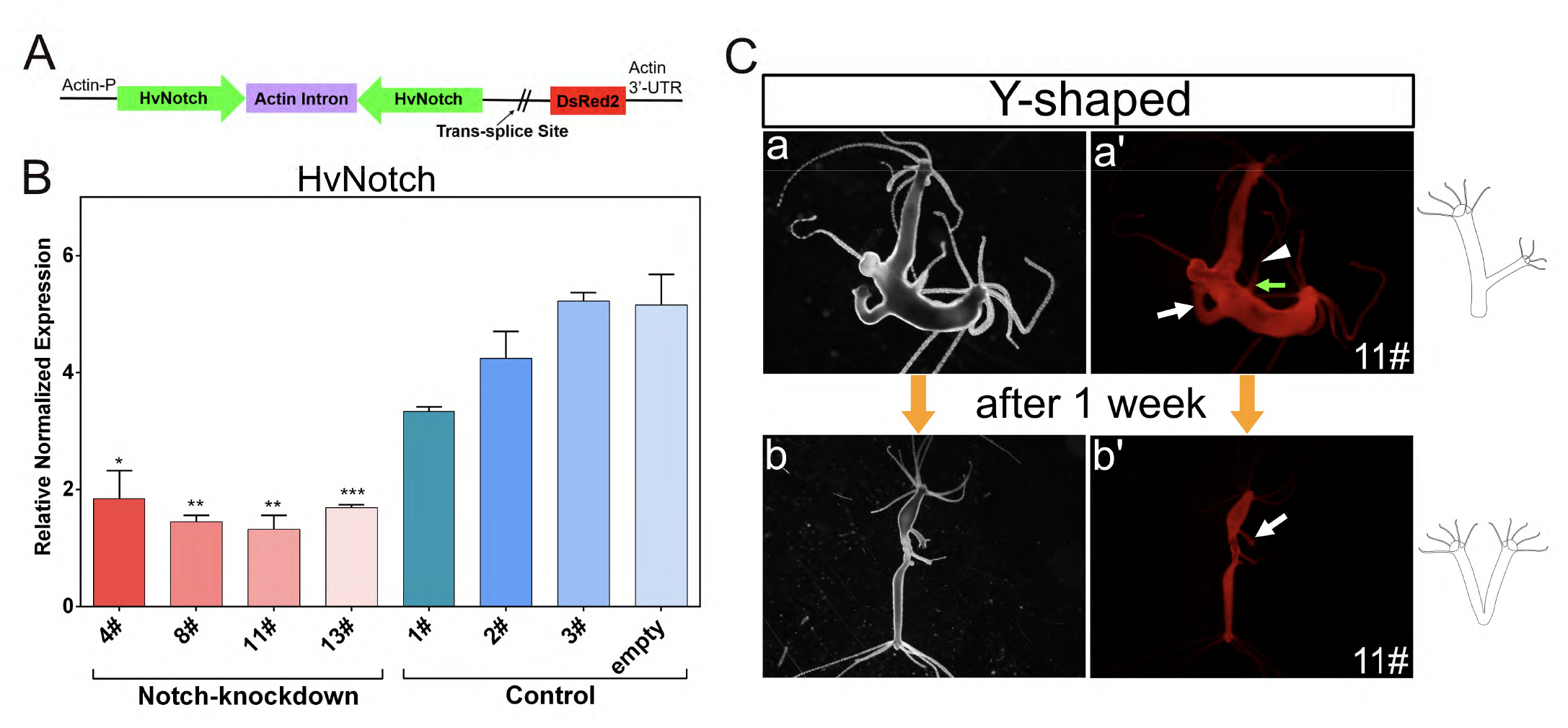
HvNotch-knockdown transgenic *Hydra*. Vector pHyVec12 used for constitutive knockdown of HvNotch. (A) The Hairpin structure containing the sequence of HvNotch in sense and antisense orientation is under the control of the actin promoter. DsRed in the same operon is expressed independently after the addition of a splice leader. (B) Diagram presents the relative normalized expression of HvNotch, as determined by RT-qPCR with mRNA isolated from four HvNotch-knockdown strains (4#, 8#, 11# and 13#), three control strains (1#, 2#, 3#) injected with control-pHyVec12 and one empty polyp strain without DsRed signals. HvNotch showed a significantly lower expression in Notch-knockdown strains in comparison with control groups (4#: p = 0.013; 8#: p = 0.0015; 11#: p = 0.0044; 13#: p = 0.0009). Primers were designed to amplify the extracellular domain of HvNotch sequences. (C) “Y-shaped” phenotype in strain 11# was observed in the initial stage (a, a’), the position of the joint moved into the foot region after one week (b, b’). a and b show light microscopy, a’ and b’ show DsRed fluorescence.

Furthermore, we noticed more “two-headed” polyps in which the second head positioned well above the budding zone (Fig. 5B, a-c). Later the body column of this second axis became longer and the joining point migrated down towards the foot region (Fig.5B, a’’ and b’’). We also observed “Y-shaped” animals, in which the second axis had originated in the budding zone (Fig.5C). Again, we ascribe “Y-shaped” polyps to the failure of bud detachment.

**Fig. 5.**
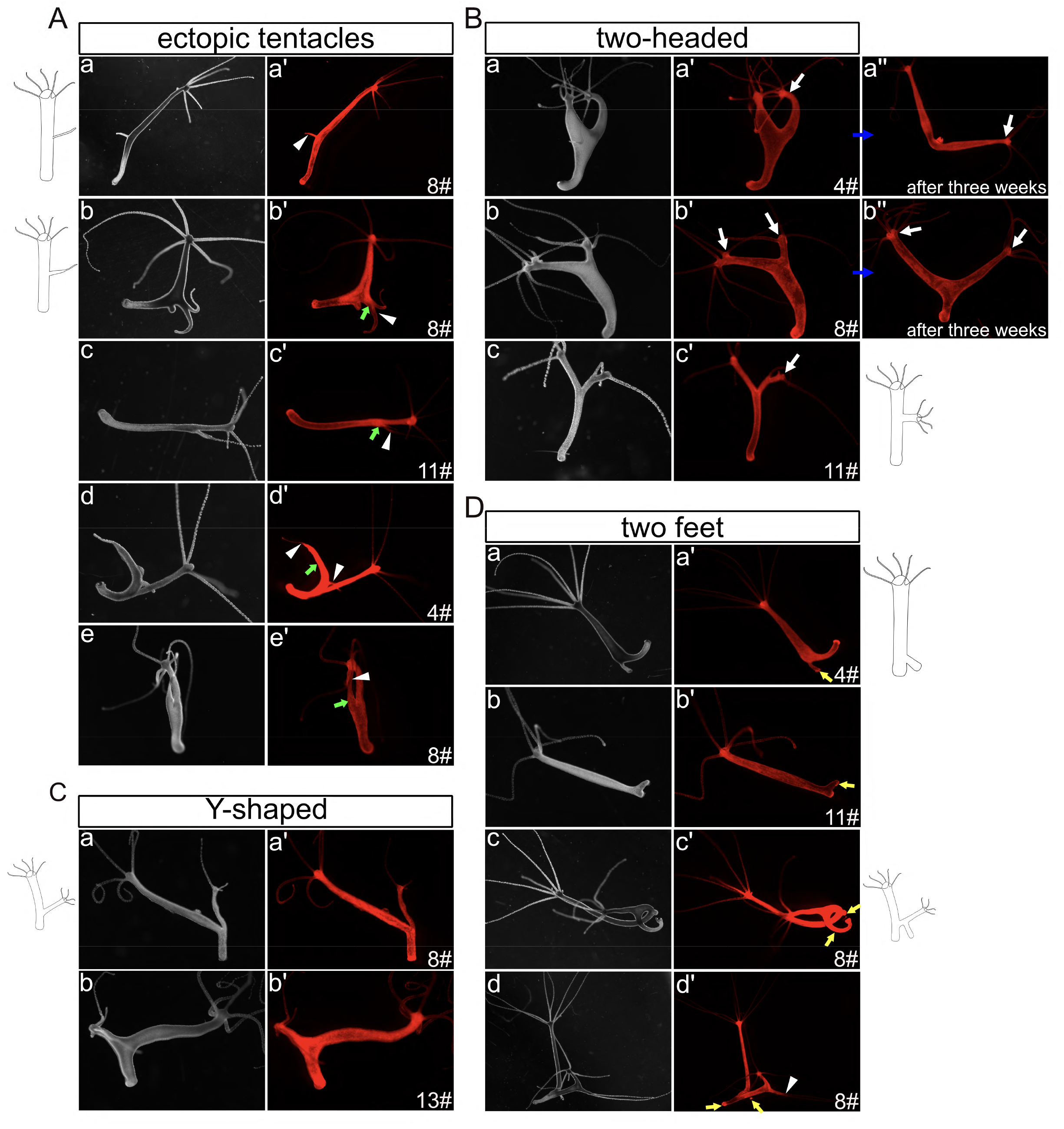
Images and drawings of Phenotypes observed in long-term Notch-knockdown transgenic *Hydra*. The images labeled with a-e represent light-microscopy, while a’-e’ display Ds-Red fluorescence. (A) “Ectopic tentacles” phenotype: (a) was similar to the phenotypes observed in HvNICD-overexpressing transgenic *Hydra*, (b-e) showed thickening of different lengths at their bases (ectopic tentacles are indicated by white triangles, thickenings at the bases of ectopic tentacles are shown by green arrows). (B) “Ectopic heads” initially located in oral half of the body column (a-c), and moving into the foot region after three weeks (a’’, b’’). Ectopic heads are indicated by white arrows. (C) “Y-shaped” polyps with the joint positioned in the budding zone. (D) “Two feet” phenotypes with extra feet indicated by yellow arrows.

**Fig. 6.**
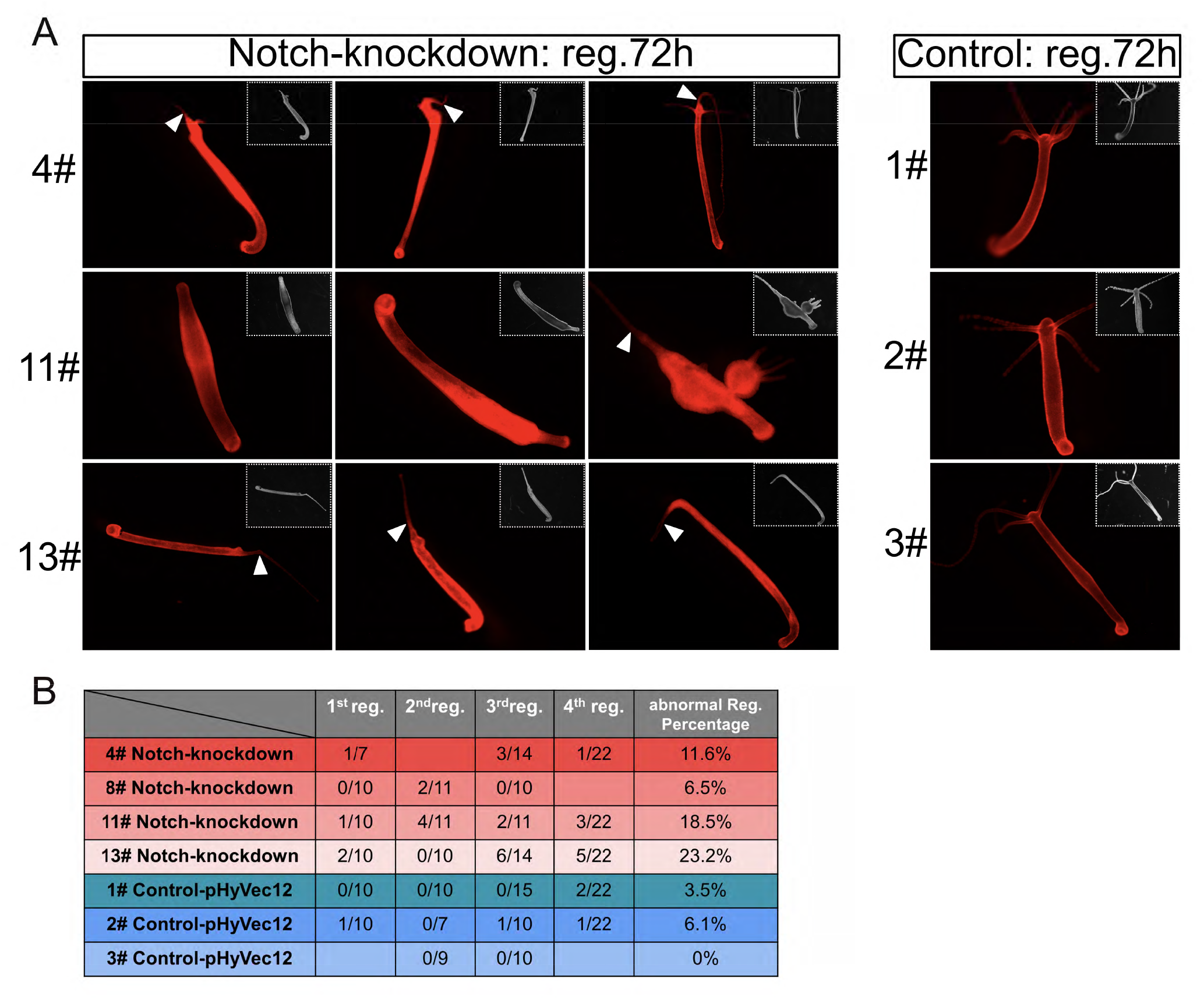
Inhibition of head regeneration by Notch-knockdown transgenic *Hydra*. (A) Images depict the DsRed fluorescence and light microscopy (inlets) of polyps from HvNotch-knockdown strains 4#, 11# and 13# 72 h after head removal, compared to control strains 1#, 2# and 3#. Abnormal regeneration involved a complete failure in regenerating head structures observed in strain 11#, or the regeneration of aberrant tentacles, as indicated by white triangles. (B) Quantification of abnormal regeneration processes in three consecutive experiments in HvNotch-knockdown and control strains

**Fig. 7.**
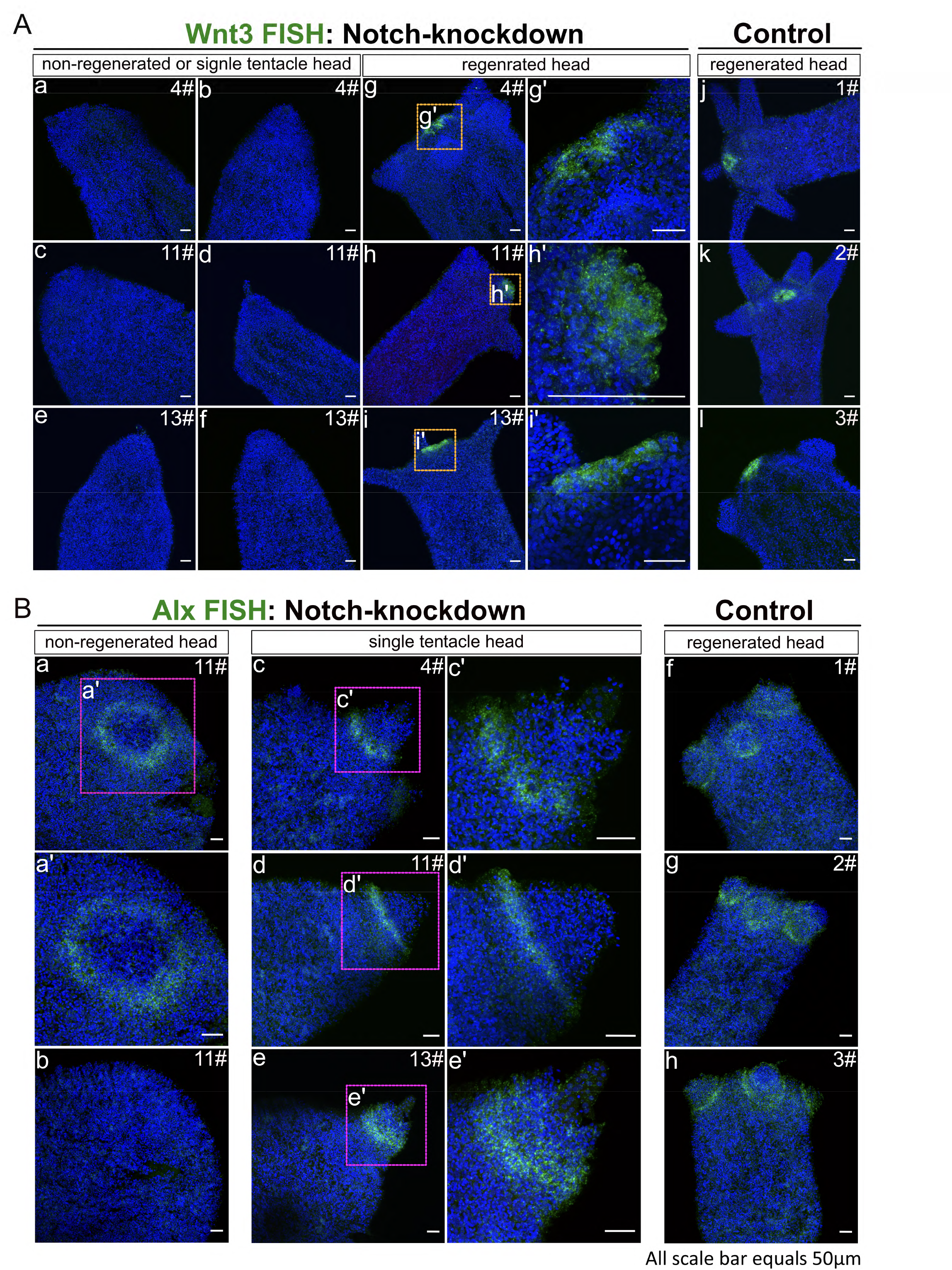
HyWnt3 and HyAlx FISH of regenerating polyps 72 h after decapitation. Stacks of laser-confocal microscopic images. (A) Expression of HyWnt3 in abnormally and normally regenerated HvNotch-knockdown polyps (a-f: non-regenerated head or single-tentacle head, g-i: normally regenerated head with enlargements g’-i’). Control polyps 1#, 2# and 3# displayed normal regeneration (j-l). (B) Expression of HyAlx in HvNotch-knockdown polyps of strain 11# with non-regenerated heads (a-b and enlargement b’), single-tentacle heads of strains 4#, 11# and 13# (c-e and enlargements c’-e’) and control polyps with normally regenerated heads of strains 1#, 2# and 3# (f-h).

A few polyps with two feet in the original polyp were detected (Fig.5D, a-b). Some “Y-shaped” animals also showed two feet (Fig.5D, c-d). In this case, the bud initial was incapable to form a foot right at the time of detachment but later developed one whilst remaining attached to the parent. Most of these phenotypes were unstable, although some simpler traits remained present after six months (supplementary Fig.S6A, B and Table S2).

### Notch-knockdown inhibited the head regeneration process

Next, we performed head regeneration experiments with Notch-knockdown polyps, choosing specimens with normal axis pattering. We found that around 20% of the 11# and the 13# strains displayed either non-regeneration or only regenerated a single tentacle. The 4# strain and 8# strains showed abnormal regeneration processes in 11% and 6% of cases, respectively (Fig.6A, 6B). However, a small percentage of control polyps also exhibited an abnormal regeneration process (Fig.6B).

We then examined the expression of HyWnt3, a marker of hypostome, and the tentacle boundary gene HyAlx in the regenerating animals three days post-decapitation, by FISH. We observed that polyps that failed to regenerate or only regenerated a single tentacle displayed a complete absence of HyWnt3 expression (Fig.7A, a-f). In contrast, normally regenerating polyps exhibited a distinct hypostomal expression pattern of HyWnt3 (Fig.7A, g-i: Notch-knockdown group; j-l: control).

For HyAlx, most non-regenerated polyps showed a large ring of HyAlx-expression at the regenerating tip (Fig.7B, a-b). In polyps with a single tentacle, HyAlx was expressed at the base of the regenerated tentacle (Fig.7B, c-e). In contrast, control polyps with fully regenerated heads displayed normal expression pattern of HyAlx at the base of each tentacle (Fig.7B, f-h). These changes in the expression patterns of HyWnt3 and HyAlx in Notch-knockdown polyps closely resembled the disturbed regeneration processes previously observed in *Hydra* polyps treated with DAPT or SAHM1^23^.

### A new model for Notch-signalling during budding in *Hydra*

The phenotypes observed in transgenic *Hydra* polyps with compromised HvNotch-expression or NICD overexpression indicate that Notch-signalling is involved in several fundamental patterning processes in *Hydra*, including budding.

To better understand these processes, we developed an initial mathematical model to illustrate the potential interaction between canonical Wnt- and Notch-signalling in *Hydra* in a simplified way. We concentrated on studying the occurrence of “Y-shaped” animals observed in budding *Hydra* treated with DAPT^24^, as well as in NICD-overexpressing and Notch-knockdown transgenic *Hydra* strains. In previous work with DAPT, we had described a sharp boundary, which is formed at the constriction stage of budding (see budding map^31^) just before foot formation. Without Notch-signalling, constriction does not occur and a foot is not formed. Even if Notch-signalling is restored later by DAPT removal, the bud does not undergo constriction and instead grows out to result in a “Y-shaped” animal. To analyse this spatio-temporal timing of Notch-induced bud constriction, we followed the ideas of Sprinzak^19,32^. We coupled a large-scale gradient in the developing bud (e.g., given by β-catenin and/or Wnt-signalling expressed at the tip of the bud) with Notch signalling by assuming a simple positive influence of canonical Wnt signalling (or another large-scale gradient apparent in the developing bud) on the Notch-ligand HyJagged, which is strongly expressed on the parent site of the boundary during the final stages of budding^22^. Furthermore, we assumed that Notch-signalling was blocked in the body-column of the parent polyp by cis-interactions between HvNotch and HyJagged. Upon simulating this system, we observed the sudden formation of a distinct ring of Notch -signalling in the most basal part of the bud, but only after the bud had reached a certain size (Fig.8A). Hence, the interplay between both systems is not only able to initiate a locally restricted ring at the future bud-foot, but also to measure the size of the protruding bud to activate HyHes-expression and following constriction at the right moment. In contrast, if Notch-signalling is inhibited in this model, bud outgrowth is not restricted, resulting in the formation of Y-shaped polyps (^24^ and this work). Interestingly, the same response occurred when we virtually overexpressed β-catenin (Fig. 8A). This is in accordance with previously reported phenotypes of transgenic *Hydra* overexpressing stabilised β-catenin, where elongated polyps were observed without any visible size limitations on the parent polyp or its buds^28^.

**Fig. 8.**
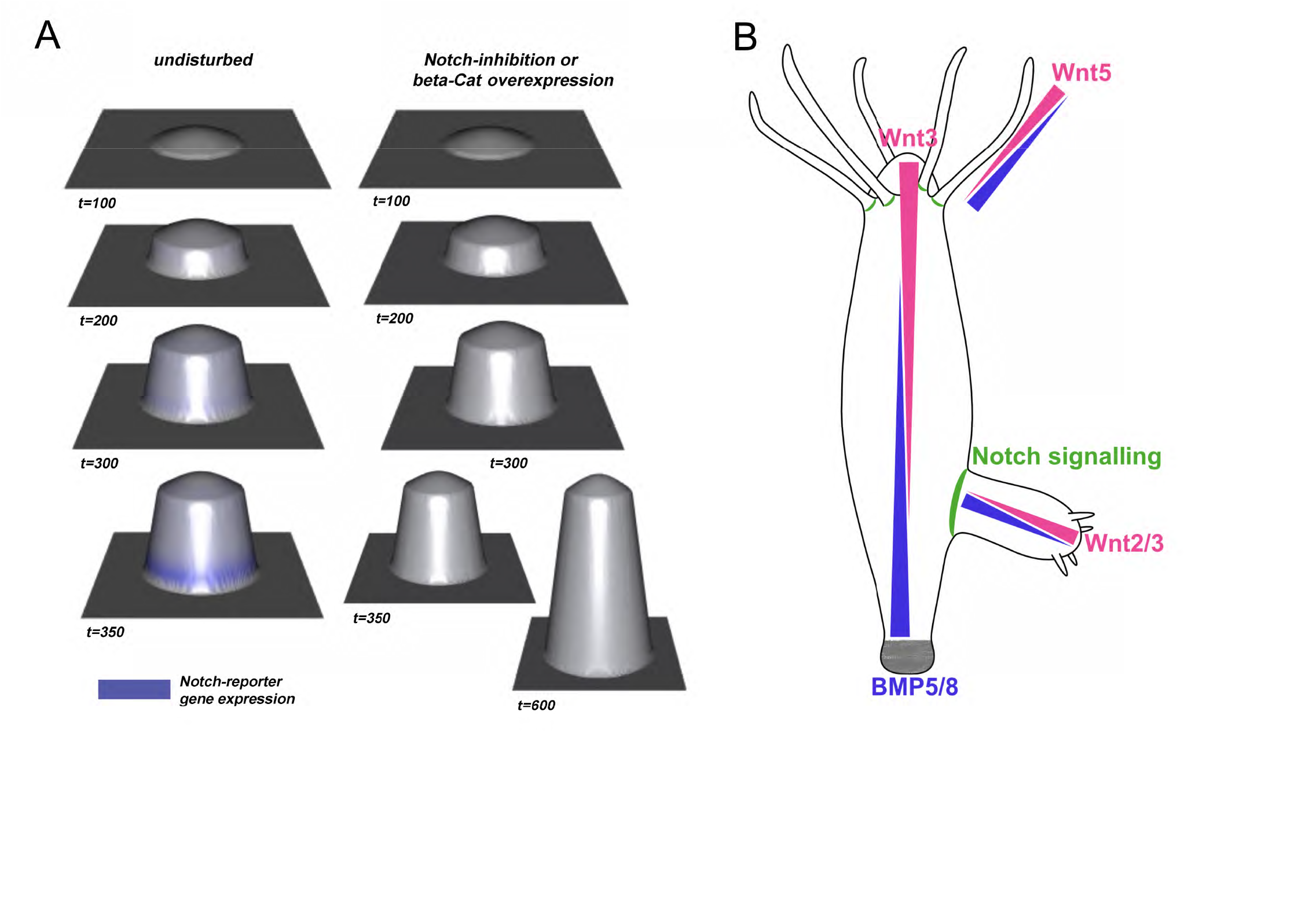
A model for the integration of Notch-signalling with long-range signalling gradients in *Hydra*. (A) Different simulated virtual time steps and phenotypes during bud-outgrowth when assuming a simple coupling between canonical Wnt- and Notch-signalling. The left-hand side shows the undisturbed system, while the right-hand side shows the impact of inhibiting Notch or overexpressing β-catenin, see also Supplementary Videos VA for undisturbed budding and VB for Notch-inhibition or β-catenin overexpression. (B) Suggested gradients of Wnt- and BMP-signalling in different parts of *Hydra*, including the body column, tentacles and buds. Gradients based on published in situ hybridisation data, Hobmeyer 2000 and Reinhardt 2004 and summarised by Meinhardt 2012. Wnt-gradient pink, BMP5-8 gradient blue; Suggested positions of Notch-signalling based on Münder 2010 and Münder 2013, green.

## Discussion

Previous work to study the function of the Notch-signalling pathway in *Hydra* was based on pharmacological pathway inhibition. It had been described that DAPT reversibly prevented nuclear translocation of NICD and similar phenotypes were obtained with a second Notch-inhibitor SAHM1, which has a completely different mode of action^21-24^. Yet, a direct proof that the observed phenotypes were solely attributable to Notch was lacking. We have now succeeded in establishing transgenic *Hydra* strains. They either expressed NICD in the whole ectoderm or endoderm, or they expressed a Notch-hairpin-RNA in both epithelial layers.

Comparison of NICD-overexpressing strains and Notch-knockdown strains revealed both similarities and important differences. Firstly, all strains showed some patterning phenotypes, such as “Y-shaped” polyps, “ectopic tentacles” and “two-headed”. Surprisingly, NICD-overexpression resulted in the down-regulation of potential Notch-target genes, including HyAlx and HySp5. In contrast, Hykayak was upregulated. These findings are consistent with the outcome of 48 h DAPT-treatment on the expression of these genes, supporting the argument of a dominant negative effect of NICD-overexpression, which has also been described in other organisms. Previous research have reported that overexpression of transgenes mostly composing of the Notch extracellular domain or the Ram23 plus Ankyrin repeat sequences led to sequestering of endogenous Notch and formation of a non-functional complex with ligands in *Drosophila*^33-35^.

In Notch-knockdown animals, the expression of HyAlx, HySp5 and Hykayak was not changed significantly. Taken together, these results suggest that NICD-overexpression had a stronger and longer-lasting effect on Notch-target genes compared to knockdown of endogenous Notch. Correspondingly, the occurrence of transgenic polyps was much lower for NICD-overexpression than for Notch-knockdown. Polyps with NICD in both epithelial layers were not obtained, whereas we established four Notch-knockdown strains. However, the similarities in the observed phenotypes can be attributed to Notch-inhibition (loss-of-function) in both cases.

NICD-overexpression revealed the presence of “Y-shaped” animals that had previously been observed after DAPT treatment. Before the bud forms its own foot, HyHes is expressed in a sharp ring of ectodermal cells at the parent-bud boundary. This process is blocked by Notch inhibition, leading to a change in the expression pattern of the *Hydra* FGF-homolog *Kringelchen*^36^. *Kringelchen* now appears in a diffused and broad zone at the base of the bud, covering both parent and bud tissue, rather than in a narrow and sharp band directly adjacent to the HyHes-expressing cells on the side of the parent^24^. This change prevents foot cell differentiation and bud detachment, resulting in the formation of “Y-shaped” animals^24^.

We suggest that NICD-overexpression in our transgenic animals inhibits ectodermal HyHes-expression when it is required to establish the parent-bud boundary. As we did not detect a down-regulation of HyHes by RT-qPCR, we furthermore propose that the high level of HyHes expression observed in the whole endoderm^37^ of *Hydra* is controlled by other factors in addition to Notch-signalling. This means that HyHes is regulated by Notch mainly in a context dependent manner, such as establishing the parent-bud boundary.

In addition to “Y-animals”, we also discovered patterning defects in NICD-overexpressing animals that were not observed with DAPT or SAHM1. These included ectopic tentacles and two- or multi-headed polyps. They reminded of phenotypes seen in animals treated with alsterpaullone or transgenic *Hydra* with overexpression of stabilised β-catenin^14,28^, both suggesting an increase of nuclear β-catenin along the body column. Occasionally, multiple heads appeared in a “bouquet” form, which were similar to those previously reported in Sp5-siRNA polyps^29,38^, which would be consistent with the downregulation of Sp5-levels found in NICD-overexpressing *Hydra*. However, the ectopic heads and tentacles on the body column indicated a change in the head activation gradient. This was also observed in Notch-knockdown *Hydra*.

All four Notch-knockdown-strains were fully transgenic in both the ectoderm and endoderm. They displayed only 30% to 50% expression of HvNotch as compared to the average of the control groups. However, the effect on the expression of Notch target genes here was much lower than in NICD-overexpressing animals. Nevertheless, we did observe some phenotypes in these strains, including “Y-shaped” animals. Moreover, Notch-knockdown affected the heads of the buds, which often displayed irregular tentacle patterns without hypostomes, resembling polyps that had undergone DAPT treatment for several shorter intervals over a longer period of time during consecutive budding processes^24^. In addition, some aberrant head structures appeared at ectopic sites in Notch-knockdown animals. They were very similar to “ectopic tentacles”, but not identical, they rather looked like heads that could only make a tentacle.

Furthermore, Notch-knockdown strains displayed a failure in head regeneration, as observed in the animals treated with DAPT^23^. In our strains, in around 20 % of cases, head regeneration either failed completely, or single tentacles were formed without hypostomes. Upon analyzing the gene expression in these regenerates, we discovered the absence of HyWnt3 when regeneration did not occur. This finding was similar to DAPT or SAHM1 treated regenerates. In contrast, the expression of the tentacle gene HyAlx could be detected in the regenerates of all knockdown animals, albeit not always in the expected ring pattern around the base of developing tentacles. Consequently, regenerates without any tentacles or with a single tentacle expressed HyAlx in a single and often broad ring. This indicated that Notch-knockdown led to a reduction of Wnt-expression during head regeneration, while not stopping HyAlx expression. As a result, a proper head could not be formed, only aberrant tentacles appeared occasionally. We explain this with the lack of a lateral inhibition process mediated by HvNotch, which is required for *Hydra* head regeneration to allow the accumulation of Wnt-3 expression at the future hypostome and to shift the expression zone of HyAlx to the base of the tentacles. When Notch is missing, the default fate of the regenerating tip is a tentacle fate. However, the lack of HyWnt3 prevents organizer formation and orderly arrangement of tentacles in this case. This idea had been described in our previous investigations on the effects of DAPT and SAHM 1 on *Hydra* head regeneration^23^.

In contrast to Notch-knockdown animals, NICD-overexpressing animals regenerated normally (supplementary Fig. S3). Moreover, they did not show aberrant head structures. We attribute this to the fact that NICD was not overexpressed in both epithelial layers, in contrast to the Notch-hairpin-RNA.

As described for NICD-overexpressing strains, Notch-knockdown strains also exhibited “two-headed” and “ectopic tentacles” phenotypes, as had been observed in animals overexpressing stabilised β-catenin and animals treated with alsterpaullone. Moreover, the knockdown animals sometimes developed two feet, which has not been seen in animals treated with DAPT or SAHM1. This constitutes a newly discovered function of Notch signalling in *Hydra*, suggesting its potential involvement in establishing or stabilising a head and/or foot activation gradient.

The effect of Notch-signalling on *Hydra* head and/or foot activation gradients is not completely unexpected given our previous findings about Notch-target genes in *Hydra*. We had described that epithelial cell genes expressed in the foot were upregulated after 48 h of DAPT treatment, including the BMP pathway component TGF-4 and APCDD1 (a negative regulator of the Wnt signalling pathway)^27^, which could explain the emergence of two-feet phenotype in Notch-knockdown animals. In contrast, head organizer genes including HyWnt7 and the transcription factor HyTCF were downregulated upon DAPT treatment. These changes have the potential to shift the head activation gradient towards the aboral end, which could explain the formation of ectopic heads above the budding zone in NICD-overexpressing and Notch-knockdown transgenic *Hydra*.

Hans Meinhardt has provided a summarised model for *Hydra* patterning in 2012, which suggested the existence of two opposing gradients of signalling molecules, one reaching from the head to the foot and the other vice versa. These two gradients are initiated by HyWnts and HyBMP5-8, respectively^11,39,40^. Moreover, these gradients are repeated in the tentacles with HyWnt5 at the tip and HyBMP5-8 at the base, and in the bud with HyWnt2 initially and later HyWnt3 at the tip and HyBMP5-8 at the base (Fig. 8B). We have extended this hypothesis by including Notch-signalling at the boundary between the parent and bud, and at the tentacle borders. In order to explain the formation of the parent-bud boundary, we have developed a mathematical model, where the gradient activity of β-catenin triggers the establishment of Notch-signals in a sharp line at this boundary. This is followed by the constriction and separation of the bud. If these signals fail to occur at the correct length of the bud, we obtain “Y-shaped” animals, indicating strict spatio-temporal requirements for this process. Within this model, the positioning of Notch-signalling depends on the concentration of Notch-receptors and ligands. When the concentrations of HvNotch and HyJagged are equal on the cell surface, they inhibit Notch-signalling in cis. The transactivation of the pathway occurs at sharp boundaries where cells with free (not cis-inhibited) Notch-receptors touch cells with free Notch ligands. Therefore, this model requires something to establish the gradients of Notch-receptors and its ligands. In *Hydra*, the HyBMP5-8/HyWnt gradients may be responsible for creating the Notch-activity gradients. In NICD-overexpressing animals, Notch-signalling could be inhibited by interactions of NICD with the endogenous HvNotch receptor on the cell membrane. In HvNotch-knockdown animals, the gradients of Notch-receptors across the length of the body column might be changed, consequently shifting the positions of the Notch-signal. In both cases, the occurrence of patterning defects can be expected.

## Methods

### Hydra culture

The injected embryos and all transgenic Hydra strains were cultured at 18 °C in Hydra medium (HM) composed of 0.29 mM CaCl_2_, 0.59 mM MgSO_4_·7H_2_O, 0.50 mM NaHCO_3_, 0.08 mM K_2_CO3. Hydra was regularly fed every two days with freshly hatched Artemia nauplii.

### Plasmid constructions

To increase the expression of NICD, 1128 base pairs of HyNotch-NICD (1648-2775 of Notch mRNA) was inserted into the vector pHyVec11 (Addgene plasmid #34794) and was driven by the Hydra actin promoter. The downstream of the NICD insert contained an intergenic sequence that leads to adding of a trans-spliced leader and the DsRed gene. For the construction of the Notch-knockdown plasmid, we designed a hairpin structure using part of the Notch-NICD sequences (1763-2283 nucleotides) in both the forward and reverse directions, separated by a 433 base pairs actin intron sequence. The entire hairpin sequence was then inserted into the vector pHyVec12 (Addgene plasmid #51851, NCBI KJ472831.1 Hydra Expression Vector pHyVec12). The Notch-Hairpin structure was under the control of actin promoter. Similar to pHyVec11, the downstream DsRed gene was connected by an intergenic region. After sequencing, the final plasmids were isolated and purified using the Qiagen Plasmid Maxi kit (QIAGEN, Cat.No.12162). Subsequently, the plasmid underwent an additional purification step with ethanol and KAc precipitation. Specifically, 10 μl of 2.5 M KAc and 250 μl of 96 % ethanol were added to a 100 μl plasmid solution obtained from the maxi-prep. The mixture was intensively mixed and incubated for 2 h at -20 °C. After incubation, the mixture was centrifuged at 15000 g for 20 min at 4°C. The resulting pellet was then washed once with 1 mL of 75 % ethanol and air-dried for 30 min to ensure the removal of any residual ethanol. Finally, we resuspended the pellet by adding 50 μL of Nuclease-free water (The resulting concentration was around 2 μg/mL).

### Generation of transgenic Hydra

The injection work was conducted in the lab of Thomas C.G. Bosch, Kiel. Following a two-week period, the embryos that were injected began to hatch. Subsequently, we screened the newly hatched polyps using a florescence microscope to identify the hatchlings with DsRed signals. Once positive stain was achieved, we fed them daily to induce the budding process and selected the buds exhibiting a higher number of transgenic signals. Throughout this procedure, we obtained some polyps that showed a notably greater concentration of signals on one side. Thus, we cut these animals and remained the pieces with more DsRed signals in order to expand the transgenic cell populations, eventually leading to the acquisition of a fully transgenic polyp.

### RT-qPCR

Total RNA was extracted using the RNeasy plus Mini kit (QIAGEN, Cat.No. A25776). The quality of the RNA was measured using the Agilent 2100 Bioanalyser with the Agilent RNA 6000 Nano kit (Agilent, Cat.No.5067-1511). Only RNA with RIN value above eight was used for cDNA synthesis using the iScript cDNA synthesis kit (Biorad, Cat.No.1708891). RT-qPCR was then performed in a 96-well plate using the PowerUp SYBR Green Master Mix (ThermoFisher, Cat.No.25742) and the CFX96^tm^ real time system from Biorad. The expression level of genes was normalized using reference genes, specifically GAPDH, EF1a and PPIB. The primer sequences used in the RT-qPCR are listed in supplementary Table S3. Statistical significance was determined based on the average value of several control groups using GraphPad Prism 6.01 with a two-tailed t-test. The corresponding P-value were expressed in the following manners. ns (no significance) for p > 0.05; * for p < 0.05; ** for p < 0.01; *** for p < 0.001; **** for p < 0.0001.

### Synthesis of RNA probes

The vector pGEM-T contains M13 primer sites in which an approximately 200 bp insert (e.g. Wnt3 and Alx) was flanked. By performing a M13 PCR, the insert was amplified and linearized, which was further verified and purified from gel with a DNA purification kit (Qiagen, Cat.No. 28704). Then, the rib probes were synthesized using SP6 or T7 polymerases together with 500 ng of M13 PCR product, DIG (digoxigenin) RNA labelling mix (Roche, Cat.No. 11277073910). According to the cloning direct of the insert, both anti-sense for the actual hybridization and sense probe for negative control were generated. Next, the DIG-labelled RNA was purification by adding 10 % of 3 M sodium acetate and 3 times of ice-cold 100 % ethanol. The labelling efficiency of probes was tested by performing a dot plot following the protocol provided by Roche. The probes that exhibited strong signals in the dot plot were considered suitable for subsequent fluorescence in-situ hybridization. The primer sequences used for amplification the insert are listed in supplementary Table S4. Approximately 25 μg of probes can be produced each cycle.

### Fluorescence in-situ hybridization

We used a protocol developed by Yashodara Abeykoon and Adrienne Cho in the laboratory of Celina Juliano. The protocol can be accessed at the following link on the protocol.io. Platform: https://www.protocols.io/view/hydra-dissociation-reaggregation-bp2l64oxdvqe/v1

### Antibody staining

Hydra were relaxed in 2 % Urethane in Hydra medium (HM) for 2 min and fixed with 4 % PFA in HM for 1 h at room temperature with gently shaking. Subsequently, the polyps were washed three times with PBS for 5 min each, and then permeabilized with 1 % Triton-X100 in PBS for 15 min. Afterwards, the polyps were blocked with a solution of 1 % BSA blocking, 0.1 % Triton-X100 in PBS for 1 h at room temperature and incubated with diluted anti-Cadherin (from Prof. Dr. Charles N. David, 1 to 1000 dilution), anti-acetylated-Tubulin (Sigma-Aldrich, Cat.No. T6793, 1 to 250 dilution) overnight at 4 °C with gentle agitation. On the next day, the polyps were washed three times with PBST (0.1 % Tween20) for 10 min each, and incubated with anti-rabbit-Alexa488, anti-mouse-Alexa649 for 2 h at room temperature. After the incubation, the polyps were washed again three times with PBST for 10 min each, nuclei were stained with DAPI (Sigma, Cat.No. D9542) at a concentration of 1 μg/ml for 15 min and mounted on slides with Vectashield anti-fade mounting medium (Biozol, Cat.No.H1000).

### Confocal imaging

A series of optical section images were captured along the Z-axis with a Leica TCS SP5-2 confocal microscope. The lasers employed in this study were Diode, Argon and Helium/Neon. After acquisition, the images were processed using ImageJ software.

DAPI nuclear staining was imaged with Diode laser with excitation wavelength at 345 nm and emission at 455 nm. Alexa488 dyes (FISH of Wnt3 and Alx, Cadherin staining) were visualized with an argon laser with excitation at 499 nm and emission at 520 nm. For Alexa 649 (acetylated-Tubulin staining), a Helium-Neon Laser with excitation at 652 nm and emission at 668 nm was used.

### Mathematical modelling of Notch signalling

The mathematical discrete model for Notch signalling during bud-outgrowth is chemically based on the MI-model as given in Sprinzak’s research^19^., where β_D represents β-catenin (or another diffusive gradient related to head identity in *Hydra*). We extend this model by coupling it to the geometrically dynamic situation of an outgrowing bud. In particular, for simulations, we discretize the unit square into 10,000 spatial pixels and define the bud-region by initially a circular region of radius 0.15 in the center of the square, slightly and spherically deformed in z-direction in order to represent the initial bud. For simulations of the bud outgrowth, starting with random initial distribution for all chemical components except β_D, on the one hand, we simulated the chemical nearest-neighbor network as given by the MI-model for 600 subsequent time steps in the bud-region only. Here, the β-catenin gradient (β_D) is prescribed in the entire domain circularly decaying from its maximum in the bud tip. On the other hand, mechano-chemical bud outgrowth is simulated by stepwise increasing the bud radius and at the same time moving bud-related pixels a constant amount (for each time step) in z-direction. This eventually leads to a stepwise protruding (slightly conical) bud shape with cells/pixels stepwise moving from the surrounding (budding-region) into the bud region, where newly augmented cells always show lower β-catenin concentrations compared to the cells augmented at the time step before.

## Acknowledgements

We want to express our gratitude to the funding agencies for supporting this work. Q.P. is funded by a CSC-grant (Chinese Scholarship Council), A.B. is funded by the German Research foundation (DFG) project BO1748-12, A.K. is supported by grants from the German Research Foundation (DFG) project KL 3475/2-1 and CRC 1461: “Neurotronics: Bio-Inspired Information Pathways”. The drawings in this document were produced using Affinity Designer 2 (Version 2.3.0) by Q.P.

## Data availability

All data presented in the main manuscript and supplementary files will be provided by the corresponding authors (Angelika Böttger and Qin Pan) upon requests.

## Author contribution statement

A.B. and Q.P. conceived this study. A.K. and J.W. conducted the induction of *Hydra* sexual reproduction, collection and injection of embryos. Q.P. established fully transgenic Hydra, performed all experiments and generated the figures. M. M. and A.M.C developed the mathematics model. Q.P. and A.B. drafted the manuscript. All authors revised and approved the manuscript.

