## Supplement for "Genetic interference with HvNotch provides new insights into the role of the Notch-signalling pathway for developmental pattern formation in *Hydra*": Supplementary files.pdf

A

| | NICD-pHyVec11 | Control-pHyVec11 |
|---|---|---|
| Injected embryos | 60 | 47 |
| Hatching Rate | 38.33%<br>(23 hatchlings) | 40.43%<br>(19 hatchling) |
| Transgenic Rate | 1.6%<br>(1 polyp) | 29.79%<br>(14 polyps) |

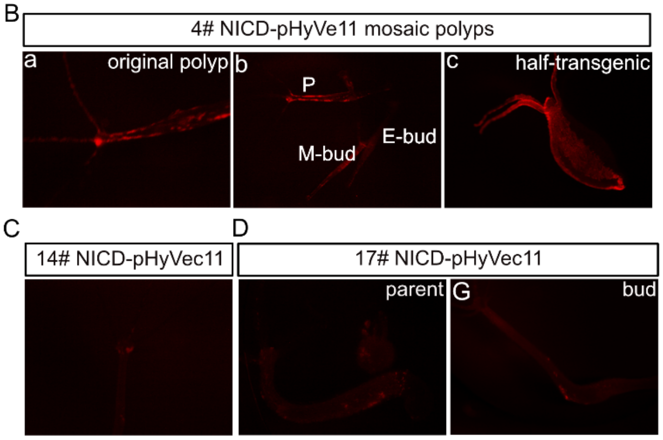


**Fig. S1 Mosaic HvNICD-overexpressing transgenic.** (A) 60 embryos were injected with HvNICD-pHyVec11, resulting in 23 hatchlings. Strain 4# had most promising transgene signals. 47 embryos were injected with control-pHyVec11, resulting in 19 hatchlings, 14 of which had good DsRed signals. (B-D) polyps from strain 4# with mosaic DsRed fluorescence. (B) 4# original polyp. (C) 4# original polyp with a developing bud and two detached buds. P: parent polyp; M-b: mosaic bud; E-b: empty bud means polyp detached from the mosaic parent, but without DsRed signals. (D) Half-transgenic polyp obtained from 4# mosaic polyp. (E) Strain 14# injected with HvNICD-pHyVec11 with few signals in the head region, migrating away from the head during development. (F-G) The original polyp and the first bud from strain 17# injected by HvNICD-pHyVec11, both with small number of scattered DsRed positive cells along the body column.

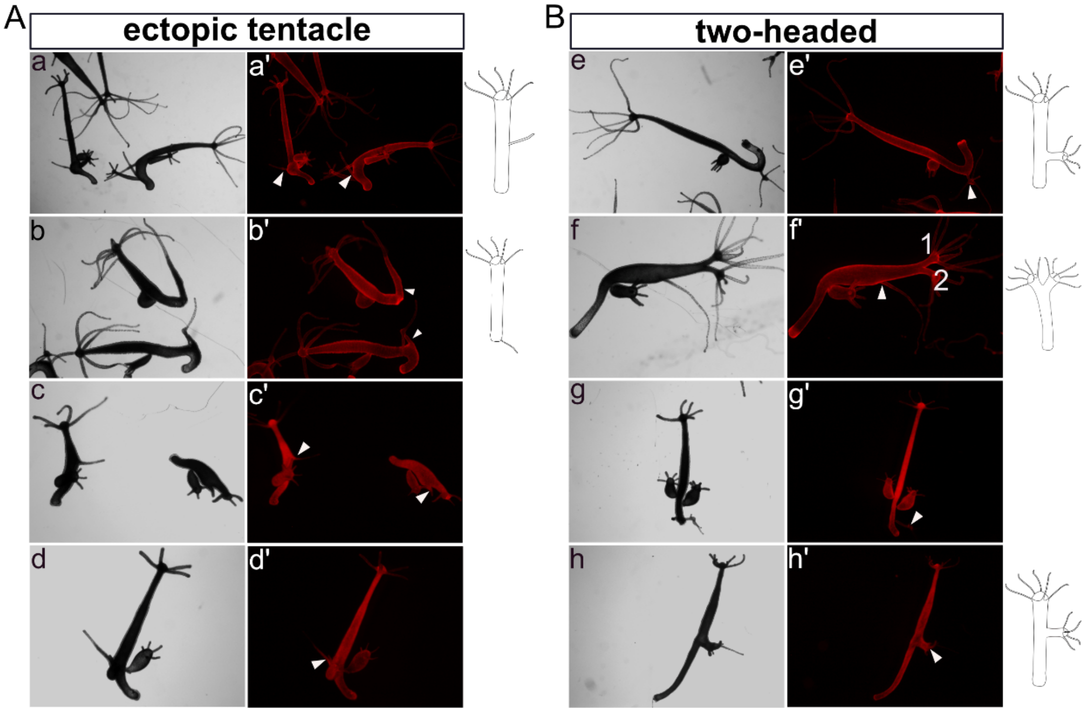


**Fig. S2 Phenotypes of HvNICD-overexpressing transgenic *Hydra* maintained after a three-week period.** (A) “Ectopic tentacles”, as indicated by white triangles in the body column (a, b ectoderm-TG; c, d endoderm-TG). (B) “two-headed” phenotype with the extra heads located in different positions along the body column and indicated with white triangles (a, b ectoderm-TG; c, d endoderm-TG).

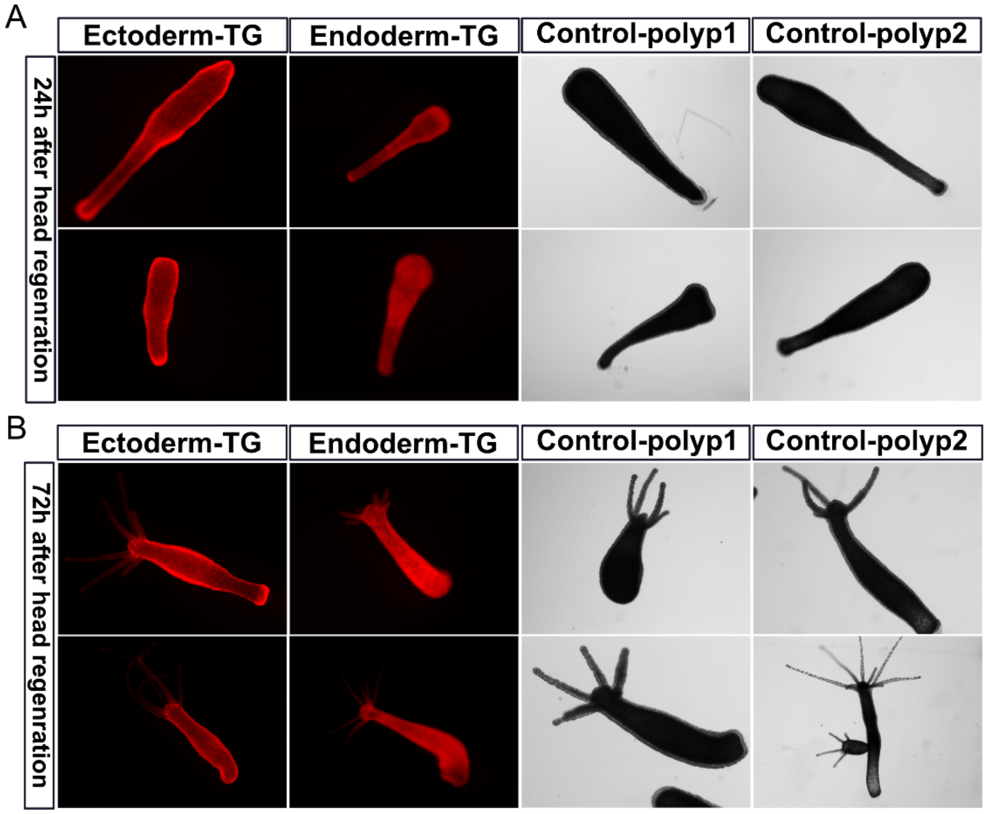


**Fig. S3 The process of head regeneration in HvNICD-overexpressing transgenic *Hydra*.** (A) The regenerating polyps from the ectoderm-TG, endoderm-TG, empty-polyp1 and empty-polyp2 24 h after decapitation. (B) 72 h after head removal, Ectoderm-TG, endoderm-TG and two control polyps all had regular head regeneration.

A

| | Notch-Hairpin-pHyVec12 | Control-pHyVec12 |
|---|---|---|
| Injected embryos | 59 | 61 |
| Hatching Rate | 38.98% (23 hatchlings) | 40.98% (25 hatchlings) |
| Transgenic Rate | 15% (9 polyps) | 16% (10 polyps) |
| Fully transgenic strains | 4#, 8#, 11#, 13# | 1#, 2#, 3# |

B

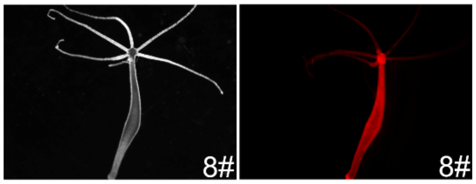


C

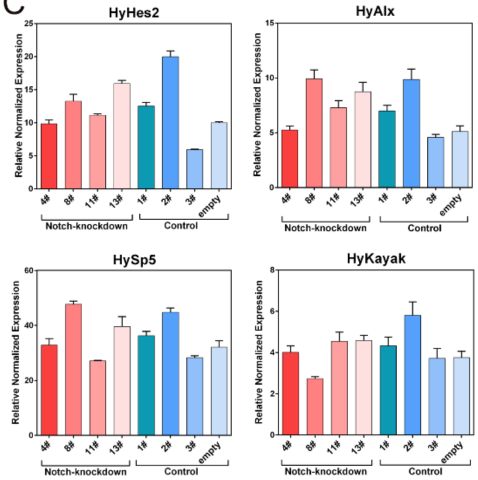


**Fig. S4 HvNotch-knockdown transgenic *Hydra* and the expression of HvNotch-target genes.** (A) 59 embryos injected with HvNotch-hairpin-pHyVec12, which produced 23 hatchlings. Among them, 9 polyps exhibited mosaic DsRed signals. In the end, 4 strains (4#, 8#, 11# and 13#) with fully transgenic signals were obtained. 61 embryos were injected with control-pHyVec12, resulting in 25 hatchling and 10 polyps with DsRed signals. From this, we generated 3 fully transgenic control strains (1#, 2# and 3#). (B) Images of fully transgenic *Hydra* from strain 8# expressing DsRed signals in both epithelial layers. (C) Diagram represents the relative normalized expression of HvNotch-target genes after RT-qPCR with mRNA from indicated Notch-knockdown and control polyps. Data for HyHes2, HyAlx, HySp5 and HyKayak are shown, differences are not substantial.

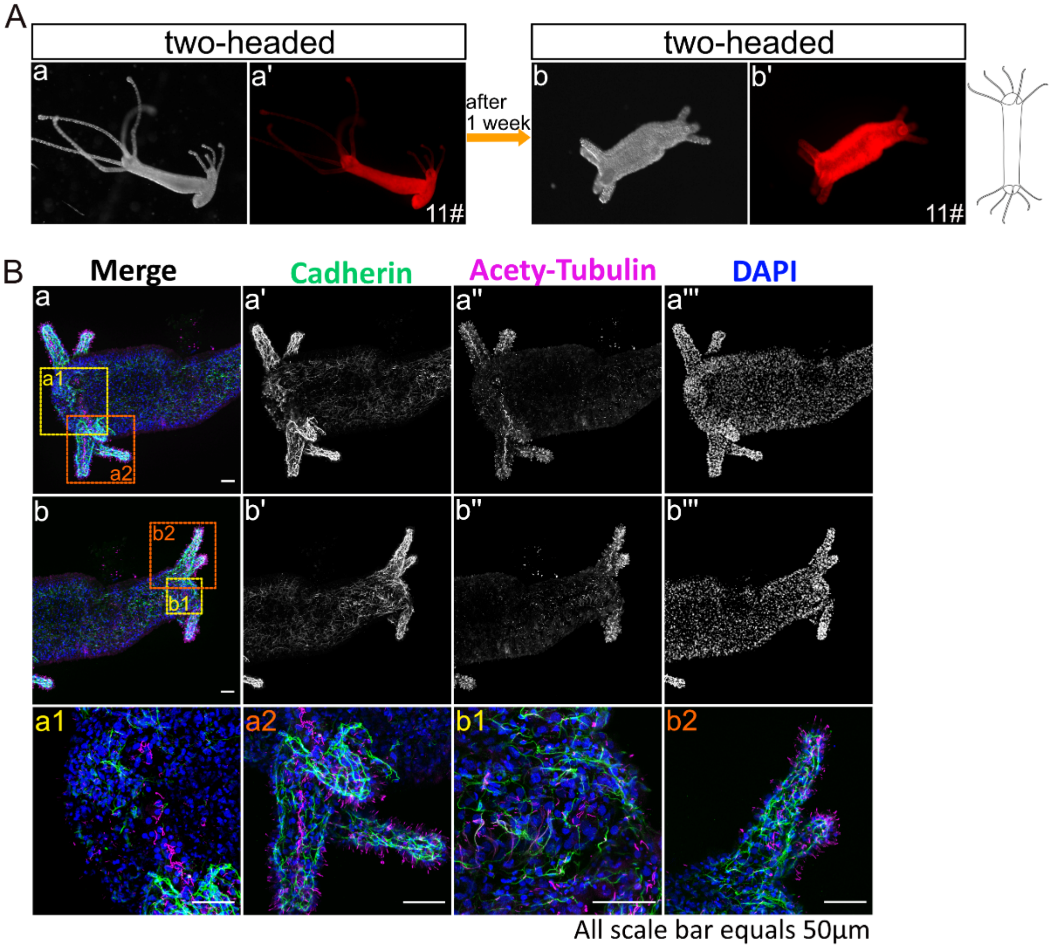


**Fig. S5 A two-headed polyp from 11# strain of HvNotch-knockdown in the initial stage.** (A) The development of this two-headed polyp with light-microscopy (a, b) and DsRed-fluorescence images (a', b') taken at the initial stage and after one-week. (B) Microscopic images of this two-headed polyp produced using confocal laser scanning after co-staining with anti-cadherin antibody to label nerve cells (black and white images a' and b'), anti-acetylated-tubulin antibodies to label cilia of nematocytes (black and white images a'' and b'') and DAPI for staining of DNA (a''' and b'''); merged images of the left head with the enlargements labeled as a1 and a2, b: merged images of the right head with enlargements labeled as b1 and b2).

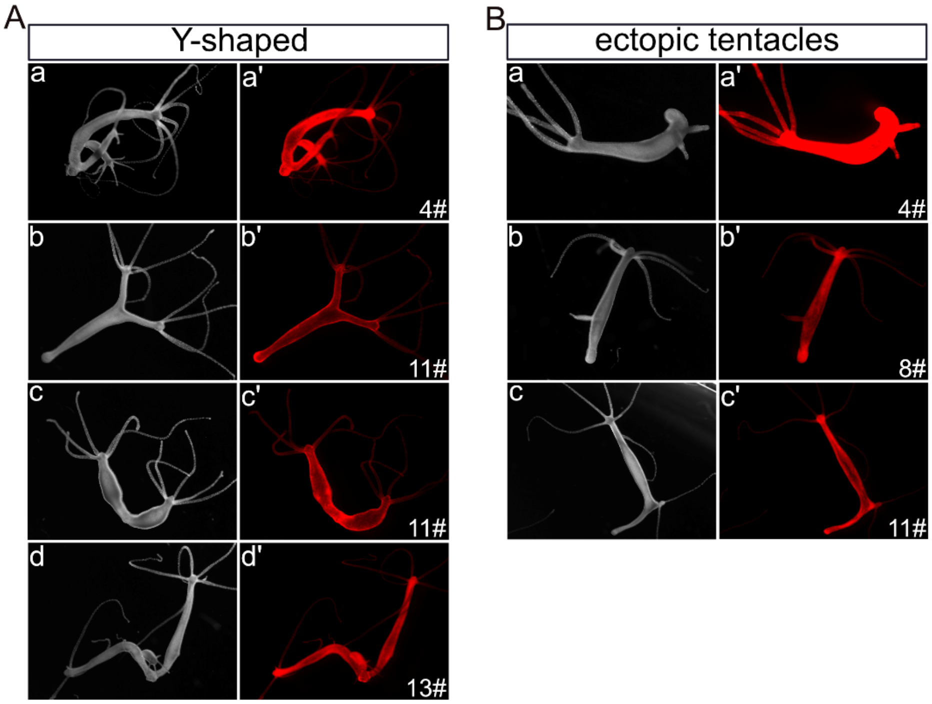


**Fig. S6 The phenotypes observed in HvNotch-knockdown transgenic *Hydra* after six months**. The images labeled with a-d represent light-microscopy, while a'-d' display Ds-Red fluorescence. (A) "Y-shaped" polyps in strains 4#, 11# and 13#. Most of joining points located in the foot region (a, c, d) while one positioned in the oral half of the body column (b). (B) Strains 4#, 8# and 11# exhibited "ectopic tentacle" phenotype with one or two ectopic tentacles in the body column.

Table S1

| Transgenic *Hydra* | Phenotypes | the 1st stage | the 2nd stage | the 3rd stage |
|---|---|---|---|---|
| Ectoderm-TG (102 polyps) | ectopic tentacles | 6 | 10 | 5 |
| | two-headed | 5 | 2 | 2 |
| | multi-headed | 0 | 3 | 1 |
| | Y-shaped | 0 | 8 | 5 |
| | combination | 0 | 4 | 2 |
| Endoderm-TG (94 polyps) | ectopic tentacles | 2 | 7 | 0 |
| | two-headed | 0 | 3 | 1 |
| | multi-headed | 0 | 1 | 1 |
| | Y-shaped | 0 | 2 | 2 |
| | combination | 0 | 0 | 1 |

Table S2

| | the 1st stage (one month) | | | |
|---|---|---|---|---|
| Transgenic Hydra | ectopic tentacles | two-headed | Y-shaped | two-feet |
| 4# Notch-knockdown | 0 | 0 | 0 | 0 |
| 8# Notch-knockdown | 0 | 1 | 0 | 0 |
| 11# Notch-knockdown | 1 | 1 | 2 | 0 |
| 13# Notch-knockdown | 0 | 0 | 0 | 0 |
| | the 2nd stage (three month) | | | |
| Transgenic *Hydra* | ectopic tentacles | two-headed | Y-shaped | two-feet |
| 4# Notch-knockdown | 2 | 1 | 3 | 1 |
| 8# Notch-knockdown | 7 | 4 | 7 | 3 |
| 11# Notch-knockdown | 5 | 3 | 6 | 1 |
| 13# Notch-knockdown | 0 | 0 | 0 | 0 |
| | the 3nd stage (three month) | | | |
| Transgenic *Hydra* | ectopic tentacles | two-headed | Y-shaped | two-feet |
| 4# Notch-knockdown | 1 | 0 | 1 | 0 |
| 8# Notch-knockdown | 1 | 0 | 0 | 0 |
| 11# Notch-knockdown | 0 | 2 | 1 | 0 |
| 13# Notch-knockdown | 0 | 0 | 2 | 0 |

**Table S1 A summary of observed phenotypes in HvNICD-overexpressing transgenic *Hydra*.** Numbers of polyps with described phenotypes at three stages (observation time points) are given for Ectoderm-TGs and Endoderm-TGs.

**Table S2 A summary of observed phenotypes in HvNotch-knockdown transgenic *Hydra*.** Numbers of polyps with described phenotypes of strains 4#, 8#, 11# and 13# of HvNotch-knockdown transgenic *Hydra* at indicated observation time points are summarized.

Table S3

| RT-qPCR primer | |
|---|---|
| NICD-P1-Fw | CAAGGCGTCATTTCTGTCAA |
| NICD-P1-Rev | ATTGGCATCAAATCCACGAT |
| NICD-P2-Fw | TAGAGGTGTTGGATGCACAAG |
| HyHes2 | GGAACTGATTCCTCGCAGAAC |
| HvNotch-Fw | TCATCATCTGACAGTGCTTT |
| HvNotch-Rev | TAGCTTGCAGCAACTTTAGG |
| HyHes-Fw | TGACGGACACAGAAAGACATC |
| HyHes-Rev | TGTCGTTTAGACTGTTGTTTATGC |
| HyAlx-Fw | GCTCGAGTACAGGTGTGGTT |
| HyAlx-Rev | AGCCGAACTACATACTGAGTTACT |
| HySP5-Fw | CGTTGCAACCCGAAGATGTC |
| HySP5-Rv | TCCGCACCCTGGAATATGAC |
| HyKayak-Fw | AACAAGTTGGCTGCTAGAAGATG |
| HyKayak-Rev | CATGGTTGTCGTGTTCAATGC |
| HyGAPDH-FW | GACAACCATTCATGCCACAA |
| HyGAPDH-FW | ACAGCTTTTGCAGCTCCAGT |
| HyEF1α-Fw | GGTCAAACCAGAGAACATGC |
| HyEF1α-Rev | TTCGCTGTATGGTGGTTCAG |
| HyPPIB-Fw | ACTGGTAAGGGAATTCTATCCA |
| HyPPIB-Rev | TACCATCCAACCATGGAGTT |

Table S4

| Cloning primers for probes of FISH | |
|---|---|
| HyAlx-Fw | TCGATTCAACTCTCCCATTTCATC |
| HyAlx-Rev | AAGGTCCGTATAGCGTCGATT |
| HyWnt3-Fw | TATCTGCGGGAGTTGCGTTT |
| HyWnt3-Rv | ACAGGTGTATTCAGGCGTCAT |

**Table S3 The list of primer sequences used in the RT-qPCR method.**

**Table S4 The list of primer sequences used to amplify the FISH-probes.**
